# Structural diversity across arbuscular mycorrhizal, ectomycorrhizal, and endophytic plant—fungus networks

**DOI:** 10.1101/269563

**Authors:** Hirokazu Toju, Hirotoshi Sato, Satoshi Yamamoto, Akifumi S. Tanabe

## Abstract

**Background:** Below-ground linkage between plant and fungal communities is one of the major drivers of terrestrial ecosystem dynamics. However, we still have limited knowledge of how such plant–fungus associations vary in their community-scale properties depending on fungal functional groups and geographic locations.

**Methods:** Based on high-throughput sequencing of root-associated fungi in eight forests along the Japanese Archipelago, we performed a comparative analysis of arbuscular mycorrhizal, ectomycorrhizal, and saprotrophic/endophytic associations across a latitudinal gradient from cool-temperate to subtropical regions.

**Results:** In most of the plant–fungus networks analyzed, host–symbiont associations were significantly specialized but lacked “nested” architecture, which has been commonly reported in plant–pollinator and plant–seed disperser networks. Meanwhile, the structure of arbuscular mycorrhizal networks was differentiated from that of ectomycorrhizal and saprotrophic/endophytic networks, characterized by high connectance. Our data also suggested that geographic factors affected the organization of plant–fungus network structure. For example, the southernmost subtropical site analyzed in this study displayed lower network-level specificity of host–symbiont associations and higher (but still low) nestedness than northern localities.

**Conclusions:** Our comparative analyses suggest that arbuscular mycorrhizal, ectomycorrhizal, and saprotrophic/endophytic plant–fungus associations often lack nested network architecture, while those associations can vary, to some extent, in their community-scale properties along a latitudinal gradient. Overall, this study provides a basis for future studies that will examine how different types of plant–fungus associations collectively structure terrestrial ecosystems.

## Background

Fungi in the below-ground biosphere are key drivers of terrestrial ecosystem processes [1–4]. Mycorrhizal fungi are considered to support land plants not only by provisioning soil nitrogen and phosphorous [5, 6] but also by increasing plants’ resistance to biotic/abiotic stress [7, 8]. Pathogenic fungi in the soil affect the survival/mortality of young plants in a major way, possibly determining spatial distributions of plant species within forest/grassland ecosystems [9, 10]. Moreover, recent mycological studies have begun to examine the poorly explored diversity of endophytic fungi, which can enhance the nutritional conditions and pathogen resistance of mycorrhizal and non-mycorrhizal plant species [11–16]. Thus, terrestrial biomes consist of multiple layers of below-ground plant–fungus interactions [17]. Nonetheless, we still have limited knowledge of the structure of such complex webs of interactions, leaving major processes in below-ground ecosystems poorly explored.

In enhancing our understanding of community- or ecosystem-level processes of below-ground plant–fungus interactions, analyses on community-scale properties of such host–symbiont associations provide essential insights. For example, if a pathogenic fungal community consists mainly of species with narrow host ranges, it as a whole is expected to restrict the emergence of dominant plant species through “negative plant–soil feedback”, contributing to the maintenance of plant species diversity within an ecosystem [18–20]. In contrast, with a high proportion of mycorrhizal fungi with narrow host ranges, their specific host species, such as Pinaceae plants hosting Suillaceae ectomycorrhizal fungi [21], will dominate the plant community through positive plant–soil feedback [20, 22]. Meanwhile, endophytic and arbuscular mycorrhizal fungi with broad host ranges [23–25] may diminish such negative and positive feedback by interlinking otherwise compartmentalized ecological dynamics (but see [26]). Therefore, concomitant analyses of community-scale properties of those multiple plant–fungus associations are of particular importance in understanding how plant–soil feedbacks organize terrestrial ecological processes.

Since the application of network science to ecology and mycology, researchers have evaluated the architecture of networks that represent linkage between plant and fungal communities [27]. Those studies have shown that arbuscular mycorrhizal [28–30], ectomycorrhizal [31], and ericaceous [32] plant–fungus networks exhibit moderate or low levels of host–symbiont specificity, while they are structured to avoid overlap of host plant ranges within fungal communities. In addition, many of those plant–fungus networks [17, 31, 33] are known to lack “nested” architecture (i.e., structure of networks wherein specialist species interact with subsets of partners of generalist species [34]), which has been commonly reported in above-ground networks of plant–pollinator and plant–seed-disperser interactions [34–36] (but see [37]). However, in those previous studies, data of different types of plant−fungus networks have been collected from different geographic localities with different sampling strategy, precluding the chance of simultaneously evaluating the effects of interaction type and geographic factors. Although comparative studies of published data provide invaluable insights [27], compiled data often vary in the molecular markers used and they may differ in appropriate null model assumptions in statistically examining network topological properties.

In this study, we compared community-scale properties of arbuscular-mycorrhizal, ectomycorrhizal, and endophytic associations across eight forest sites spanning from cool-temperate to subtropical regions in Japan. Based on high-throughput sequencing of root-associated fungi, we obtained network data depicting how multiple plant species are associated with respective functional groups of fungi in each of the eight forests. We then examined how structure varied depending on categories of plant–fungus associations and geographic locations. Overall, this study provides a first step for integrating insights into community-scale properties of multiple types of below-ground plant–fungus associations and their ecosystem-level consequences.

## Methods

### Terminology

In analyzing metadata of community-scale properties of plant–fungus associations, we need to use consistent terminology that can be applied to a wide range of host–symbiont associations. While plant–fungus network properties have been compared within a single functional group of fungi (e.g., arbuscular mycorrhizal or ectomycorrhizal fungi) in most studies, we herein target not only arbuscular mycorrhizal and ectomycorrhizal fungi but also pathogenic and saprotrophic/endophytic fungi. Given that those functional groups of fungi vary considerably in their microscopic structure within plant tissue [8], developing a general criterion for mutualistic/antagonistic interactions with host plants is impossible. Thus, we targeted all the fungi detected by high-throughput sequencing and the data described below could contain not only mutualistic/antagonistic fungi but also commensalistic fungi merely adhering to plant roots [38]. In this sense, our data represented symbiotic relationships in the broad sense, i.e., intimate physical connections between organisms [17, 39].

### Sampling

We collected root samples at eight forest sites (four cool-temperate, one warm-temperate, and three subtropical forests) across the entire range of the Japanese Archipelago (45.042-24.407 °N; Fig. 1A; Additional file 1: Data S1). In each forest, 2-cm segment of terminal roots were collected from 3-cm below the soil surface at 1-m horizontal intervals. In each forest, 383 terminal root samples were collected. Those roots were collected indiscriminately regarding root morphology or apparent mycorrhizal type so that the samples as a whole represented the relative frequency of plant–fungal associations in the horizon in each forest [40]. Therefore, while the sample sets consisted mainly of woody plants, they also included herbaceous plants (Additional file 2: Data S2). Each root sample was preserved in 70% ethanol and stored at -25 °C until DNA extraction.

**Fig. 1.**
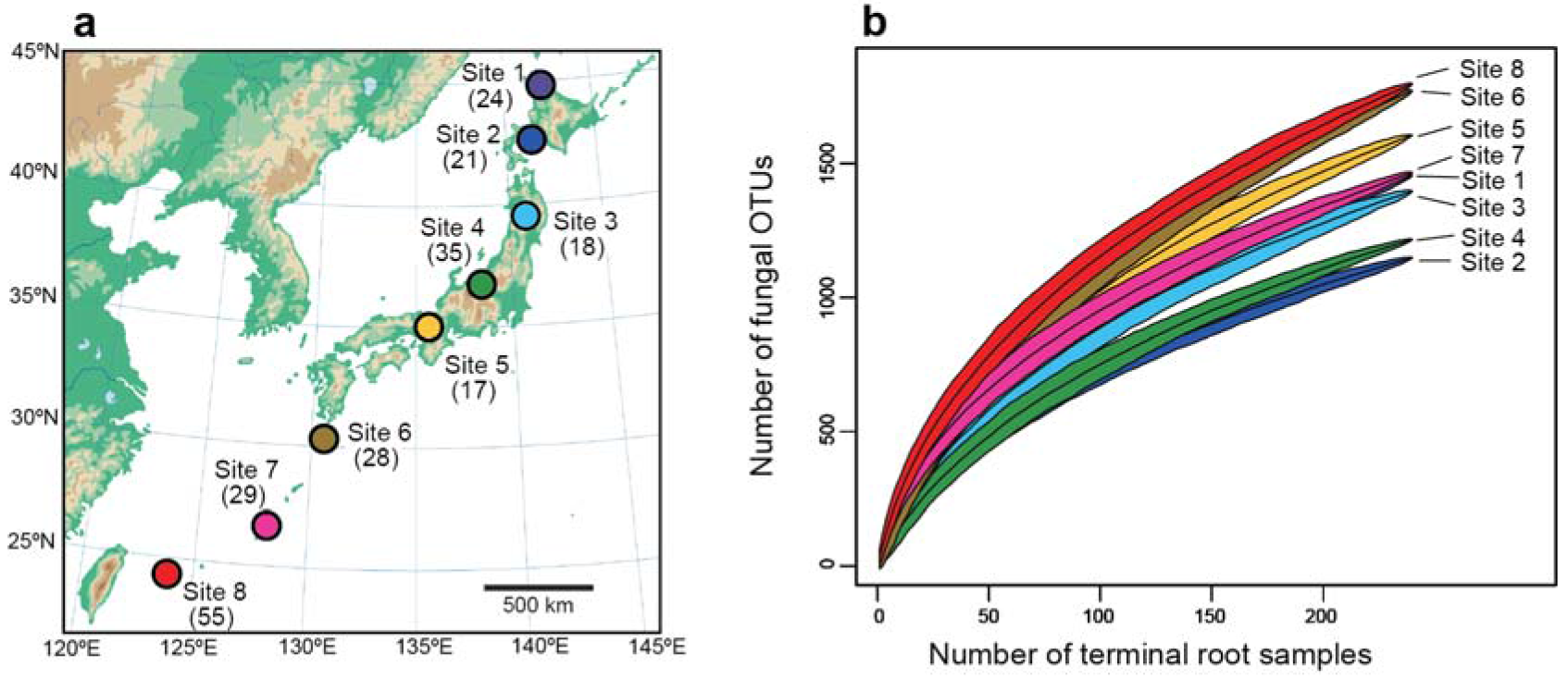
Study sites. **a** Map of study sites. In each forest site, a number in a parenthesis indicates the number of plant species/taxa observed in the 240 terminal root samples from which sequencing data were successfully obtained. **b** Relationship between the number of samples and that of plant species/taxa observed. A rarefaction curve obtained from 240 terminal-root samples is shown for each study site.

### Molecular analysis

Each root sample was placed in 70% ethanol with 1-mm zirconium balls and it was then shaken at 15 Hz for 2 min with a TissueLyser II (Qiagen) [23]. The washed roots were subsequently pulverized by shaking with 4-mm zirconium balls at 25 Hz for 3 min. DNA extraction was then performed with a cetyltrimethylammonium bromide method [41].

The internal transcribed spacer 1 (ITS1) region of root-associated fungi was amplified with the primers ITS1F_KYO1 and ITS2_KYO2, which target not only Ascomycota and Basidiomycota fungi but also diverse non-Dikarya (e.g., Glomeromycota) taxa [42]. We used the forward primer ITS1F-KYO1 fused with 3—6-mer Ns for improved Illumina sequencing quality [43] and the forward Illumina sequencing primer (5’- TCG TCG GCA GCG TCA GAT GTG TAT AAG AGA CAG-[3—6-mer Ns] - [ITS1-KYO2] -3’) and the reverse primer ITS2-KYO2 fused with 3—6-mer Ns and the reverse sequencing primer (5’- GTC TCG TGG GCT CGG AGA TGT GTA TAA GAG ACA G [3—6-mer Ns] — [ITS2_KYO2] -3’). The DNA polymerase system of KOD FX Neo (Toyobo) was used with a temperature profile of 94 °C for 2 min, followed by 35 cycles at 98 °C for 10 s, 50 °C for 30 s, 68 °C for 50 s, and a final extension at 68 °C for 5 min. The ramp rate was set to 1 °C/sec to prevent the generation of chimeric sequences [44]. Illumina sequencing adaptors were then added to each sample in the subsequent PCR using the forward primers consisting of the P5 Illumina adaptor, 8-mer tags for sample identification [45], and a partial sequence of the sequencing primer (5’- AAT GAT ACG GCG ACC ACC GAG ATC TAC AC - [8-mer index] - TCG TCG GCA GCG TC -3’) and the reverse primers consisting of the P7 adaptor, 8-mer tags, and a partial sequence of the sequencing primer (5’- CAA GCA GAA GAC GGC ATA CGA GAT - [8-mer index] - GTC TCG TGG GCT CGG -35’). In the reaction, KOD FX Neo was used with a temperature profile of 94 °C for 2 min, followed by 8 cycles at 98 °C for 10 s, 55 °C for 30 s, 68 °C for 50 s, and a final extension at 68 °C for 5 min. The PCR amplicons of 384 samples in each forest (including one PCR negative control) were pooled with equal volume after a purification/equalization process with AMPureXP Kit (Beckman Coulter).

For the identification of plants, another set of PCR was performed targeting chloroplast *rbcL* region with rbcL_F3 and rbcL_R4 primers [40]. The fusion primer design, DNA polymerase system, temperature profiles, and purification processes used in the *rbcL* analysis were the same as those of the fungal ITS analysis. The ITS and *rbcL* libraries were processed in two Illumina MiSeq runs, in each of which samples of four forest sites were combined (run center: KYOTO-HE) (2 × 250 cycles; 15% PhiX spike-in).

### Bioinformatics

In total, 17,724,456 and 17,228,848 reads were obtained for the first and second MiSeq runs. The raw sequencing data were converted into FASTQ files using the program bcl2fastq 1.8.4 provided by Illumina. The FASTQ files were then demultiplexed using the program Claident v0.2.2016.07.05 [46, 47]. To avoid possible errors resulting from low-quality index sequences, the sequencing reads whose 8-mer index positions included nucleotides with low (< 30) quality scores were discarded in this process. As reverse sequences output by Illumina sequencers have lower quality values than forward sequences, we used only forward sequences after removing low-quality 3’-ends using Claident (sequencing data deposit: DDBJ DRA accession: DRA006339). Noisy reads were subsequently discarded and the reads that passed the filtering process were clustered using VSEARCH [48] as implemented in Claident. The threshold sequencing similarities in the clustering were set to 97% for fungal ITS and 98% for *rbcL*, respectively. While sequence similarity values have been set to 97% in most ITS analyses of Ascomycota and Basidiomycota fungi [49] (see also [50]), a recent study showed that Glomeromycota fungi generally had much higher intraspecific ITS-sequence variation than Dikarya fungi [51]. Therefore, we performed an additional clustering analysis with a 94% cutoff similarity for defining Glomeromycota OTUs. Note that changing cut-off similarities (81-97%) did not qualitatively change statistical properties of plant–fungus network structure in a previous study [17]. The taxonomic assignment of the OTUs (Additional files 3-4: Data S3-4) was conducted based on the combination of the query-centric auto-*k*-nearest neighbor (QCauto) method [46] and the lowest common ancestor (LCA) algorithm [52] as implemented in Claident. Note that taxonomic identification results based on the QCauto–LCA approach were comparable to, or sometimes more accurate than, those with the alternative approach combining the UCLUST algorithm [53] with the UNITE database [54] [see [32] and [55] for detailed comparison of the QCautoLCA and UCLUST–UNITE approaches]. The functional group of each fungal OTU was inferred using the program FUNGuild 1.0 [56]. For 44.1 % (3560/8080) of fungal OTUs, functional group information was inferred (Additional file 1: Data S1).

The obtained information of *rbcL* OTUs was used to identify each root sample, although species-level taxonomic information was unavailable for some plant taxa in each forest due to the relatively low variability of the chloroplast region [57]. Thus, we also used the information of the ITS sequencing libraries, which included not only fungal but also host plant sequencing reads: there were plant taxa that could not be identified to species even with ITS information. Based on the *rbcL* and ITS information of plant sequences, possibly contaminated samples were removed from the dataset.

For each of the eight forests, we then obtained a sample (row) × fungal OTU (column) data matrix, in which a cell entry depicted the number of sequencing reads of an OTU in a sample. The cell entries whose read counts represented less than 0.1% of the total read count of each sample were subsequently excluded because those rare entries could derive from contaminations from soil or PCR/sequencing errors [58]. The filtered matrices were then rarefied to 1,000 reads per sample using the “rrarefy” function of the vegan 2.4-3 package [59] of R 3.4.1 [60]. As the number of samples with 1,000 or more reads varied among the eight forests examined (240–288 samples), it was equalized by randomly sampling 240 samples without duplication in each forest (“sample-level matrices”; Additional file 2: Data S2).

Based on the sample-level matrices, we obtained another type of matrices, in which a cell indicated the number of samples representing associations between a plant species/taxa –(row) and a fungal OTU (column) (“species-level matrices”; Additional file 5: Data S5). In addition to the matrix indicating associations between all fungal OTUs and their host plants (ALL), a series of partial network matrices representing respective fungal functional groups were obtained by selecting arbuscular mycorrhizal (AM), ectomycorrhizal (ECM), potentially pathogenic (PATHO), and saprotrophic/endophytic (SAPENDO) fungal OTUs in each forest (Additional file 6: Data S6). Due to the limited availability of information of fungal ecology, functional groups of many fungal OTUs could not be estimated and there were only 9–25 fungal OTUs inferred to be plant pathogens in respective forests (Additional files 1 and 5: Data S1 and S5).

### Data analysis

Based on the sample-level matrices, relationship between the number of samples and that of observed fungal OTUs was analyzed for each forest using the “specaccum” function of the vegan package. The community-scale plant–fungus associations represented by the species-level matrices (“ALL” network matrices; Additional file 5: Data S5) were visualized using the program GePhi 0.9.1 [61] with “ForceAtlas2” layout algorithm [62]. We then analyzed the statistical properties of the ALL networks and partial networks (Additional file 6: Data S6) in terms of the *H*_2_’ metric of network-level interaction specificity [63], which has been frequently used to measure the degree of interaction specificity in host–symbiont networks [64, 65]. The plant–fungus associations were evaluated also by the weighted NODF metric [66] of network nestedness [34], which measures the degree to which specialists (species with narrow partner ranges) interact with partners of generalists (species with broad partner ranges) in the same guild or trophic level. We further examined how host plant ranges were differentiated within the fungal community of each forest based on checkerboard scores [67]: a high/low score of the checkerboard index indicates host differentiation/overlap within a guild or trophic level [65]. Although modularity is another important index frequently used in ecological network studies [35], its computation was too time-consuming to be applied to randomization analyses (see below) of our present datasets consisting of more than 1,000 fungal OTUs and their host plants. Note that we previously found that below-ground plant–fungal associations generally showed statistically significant but low network modularity [17, 32, 65].

As estimates of network indices could vary depending on species compositions of examined communities, we standardized the indices as

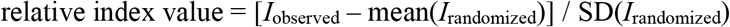

where *I*_observed_ was the index estimate of the observed data matrix, and mean(*I*_randomized_) and SD(*I*_randomized_) were the mean and standardized deviation of the index values of randomized matrices [65]. Randomized matrices were obtained by shuffling host-plant labels in the sample-level matrices and subsequently converting the randomized sample-level matrices into randomized species-level matrices. Although we used two additional methods [“r2dtable” [68] and “vaznull” [69] methods] of matrix randomization in our previous studies of plant–fungus networks [17, 65], they were too time-consuming to be used in the present large datasets: note that the three randomization methods compared in those previous studies yielded qualitatively similar results [17, 65]. The number of randomizations was set to 1,000 for *H*_2_’/nestedness analyses and 100 for checkerboard-score analyses, which required substantial computing time.

Based on the network indices, we examined how the community-scale properties of the plant–fungus associations varied among local forests and fungal functional groups. For each of interaction specificity (relative *H*_*2*_’), nestedness (relative weighted NODF nestedness), and checkerboard index (relative checkerboard values), an ANOVA model was constructed by incorporating locality (forest sites), fungal functional group, number of plant species/taxa, number of fungal OTUs, and network connectance (the proportion of non-zero entries in community matrices) as explanatory variables. The variation in the plant–fungus network properties was visualized based on a principal component analysis based on a correlation matrix: the variables included were *H*_*2*_’ interaction specificity, NODF nestedness, checkerboard index, number of plant species/taxa, number of fungal OTUs, proportion of fungal OTUs to plant species/taxa, and connectance.

## Results

Total fungal OTU richness was higher in warm-temperate and subtropical forests than in cool-temperate forests (Figs. 1b and 2). The OTU richness of AM fungi was higher in the three subtropical forests, while that of ECM fungi decreased in the southern forests (Fig. 3a). The ratio of the total number of fungal OTUs to the number of plant species/taxa varied among forests, although there was seemingly no systematic variation between cool-temperate and the other (warm temperate and subtropical) localities (Fig. 3b). Connectance varied among forests as well, while it was consistently higher in AM than in ALL, ECM, and SAPENDO networks/partial networks in seven of the eight study forests (Fig. 3c). The connectance of PATHO partial networks varied considerably among forests presumably due to low OTU richness and the resultant uncertainty in index estimation.

**Fig. 2.**
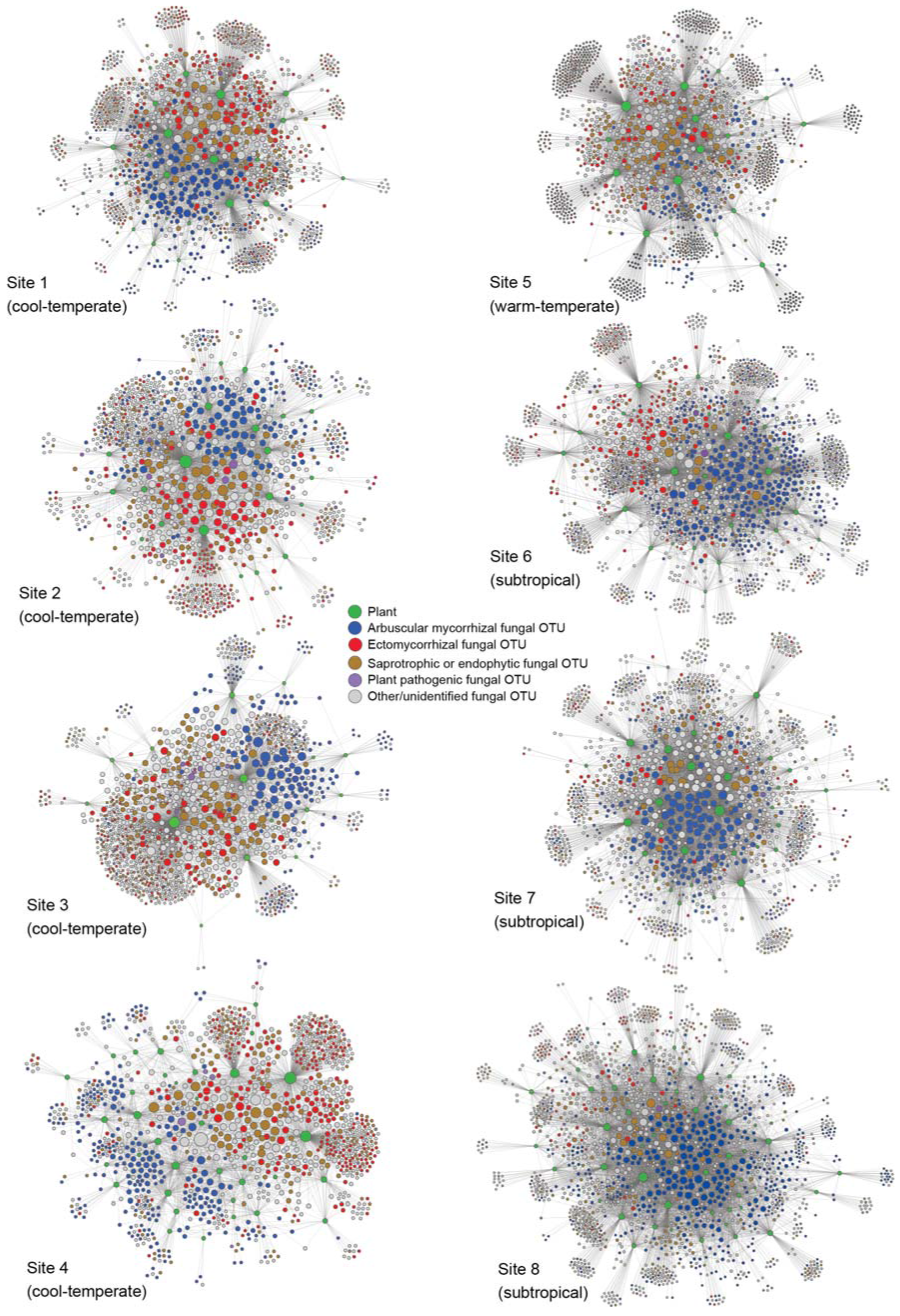
Below-ground plant–fungus networks. The “ALL” network involving all the root-associated fungal OTUs detected and their host plant species/taxa is shown for each forest. The OTUs/species in the networks are arranged with the “ForceAtlas2” layout algorithm [62]. Size of circles represents betweenness centrality scores compared within plant/fungal community.

The relative *H*_*2*_’ metric of interaction specificity significantly varied among forests but not among fungal functional groups when the effects of plant diversity, fungal OTU richness, and connectance were controlled in an ANOVA model (Table 1; Fig. 3d). The relative nestedness of the ALL matrices of plant–fungus associations was lower than zero in most forests but not in the southern most subtropical forest (Fig. 3e; Additional file 7: Data S7). Overall, plant–fungus associations in ALL networks were more specialized (Fig. 3d) and less nested (Fig. 3e) than those of partial networks. In addition, fungal OTUs in ALL networks displayed stronger differentiation of host ranges than those in partial networks (Fig. 3f).

**Fig. 3.**
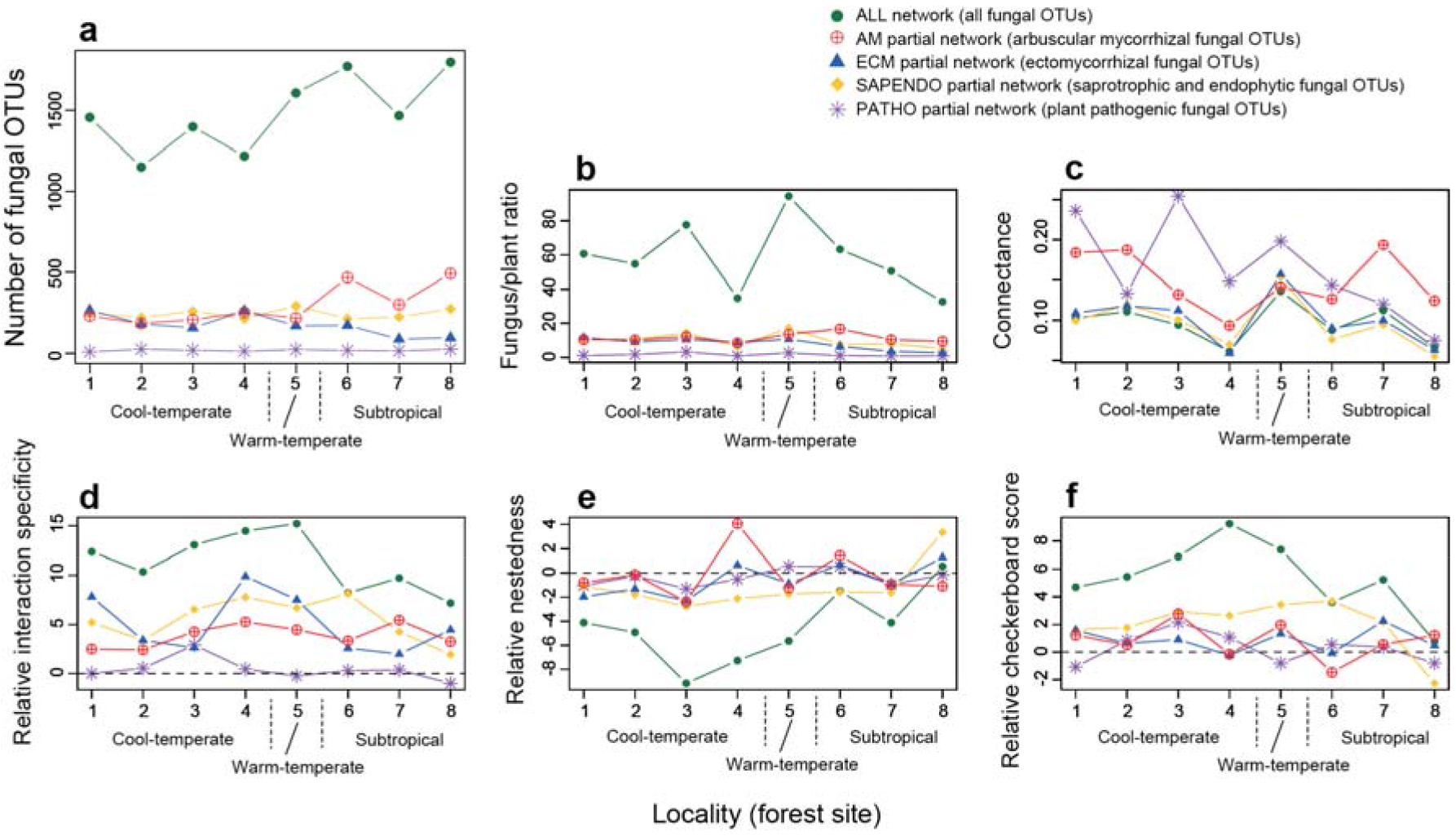
Network properties. The index scores representing the architecture of plant fungus networks/partial networks are shown across the eight forests examined. **a** The number of fungal OTUs. The code numbers of forest sites correspond to those shown in Figure 1. **b** The ratio of the number of fungal OTUs to that of the plant species/taxa involved in each network/partial network. **c** Connectance (the proportion of non-zero entries in a species-level matrix). **d** Network-level interaction specificity (relative *H*_2_’). **e** Nestedness (relative weighted NODF nestedness). **f** Host range differentiation (relative checkerboard score). For relative interaction specificity, relative nestedness, and relative checkerboard score (**d**-**f**), scores higher/lower than 2 roughly indicate that observed network index values are higher/lower than expected by chance (see Additional file 7: Data S7 for detailed results of the randomization test).

**Table 1.** Potential factors contributing to variation in plant–fungus network structure. For each response variable representing network structure, an ANOVA model including the number of plant species/taxa, that of fungal OTUs, network connectance, sampling locality, and fungal functional groups (i.e., categories of plant–fungus networks) was constructed. *P* values significant after a Bonferroni correction are shown in bold for ANOVA model.

| Response variable | Explanatory variable | df | *F* | *P* |
| --- | --- | --- | --- | --- |
| Relative interaction specificity | No. plant species/taxa | 1 | 4.6 | 0.0426 |
|  | No. fungal OTUs | 1 | 96.9 | < **0.0001** |
|  | Connectance | 1 | 10.1 | **0.0040** |
|  | Locality | 7 | 4.0 | **0.0048** |
|  | Functional group | 4 | 2.6 | 0.0572 |
| Relative nestedness | No. plant species/taxa | 1 | 5.1 | 0.0322 |
|  | No. fungal OTUs | 1 | 39.3 | < **0.0001** |
|  | Connectance | 1 | 2.4 | 0.1308 |
|  | Locality | 7 | 1.5 | 0.2216 |
|  | Functional group | 4 | 2.1 | 0.1179 |
| Relative checkerboard score | No. plant species/taxa | 1 | 1.0 | 0.3182 |
|  | No. fungal OTUs | 1 | 62.9 | < **0.0001** |
|  | Connectance | 1 | 5.6 | 0.0262 |
|  | Locality | 7 | 1.3 | 0.2772 |
|  | Functional group | 4 | 4.0 | **0.0121** |

After taking into account plant and fungal diversity in an ANOVA model, neither locality nor fungal functional group explained the variation in relative nestedness (Table 1). The relative checkerboard scores varied among localities (Fig. 3f), although the effects of locality were non-significant in an ANOVA model (Table 1). The ANOVA model showed that the variation in relative checkerboard scores was explained, to some extent, by fungal functional groups.

In the principal component analysis of network indices, ALL, PATHO and other networks/partial networks were separated by the first principal component, which represented high plant diversity, fungal OTU richness, relative *H*_*2*_’, and relative checkerboard scores as well as low relative nestedness (Fig. 4a). By incorporating the third principal component, which represented high fungal OTU richness and connectance, the cluster of AM partial networks and that of ECM and SAPENDO partial networks were grouped with some overlap (Fig. 4b).

**Fig. 4.**
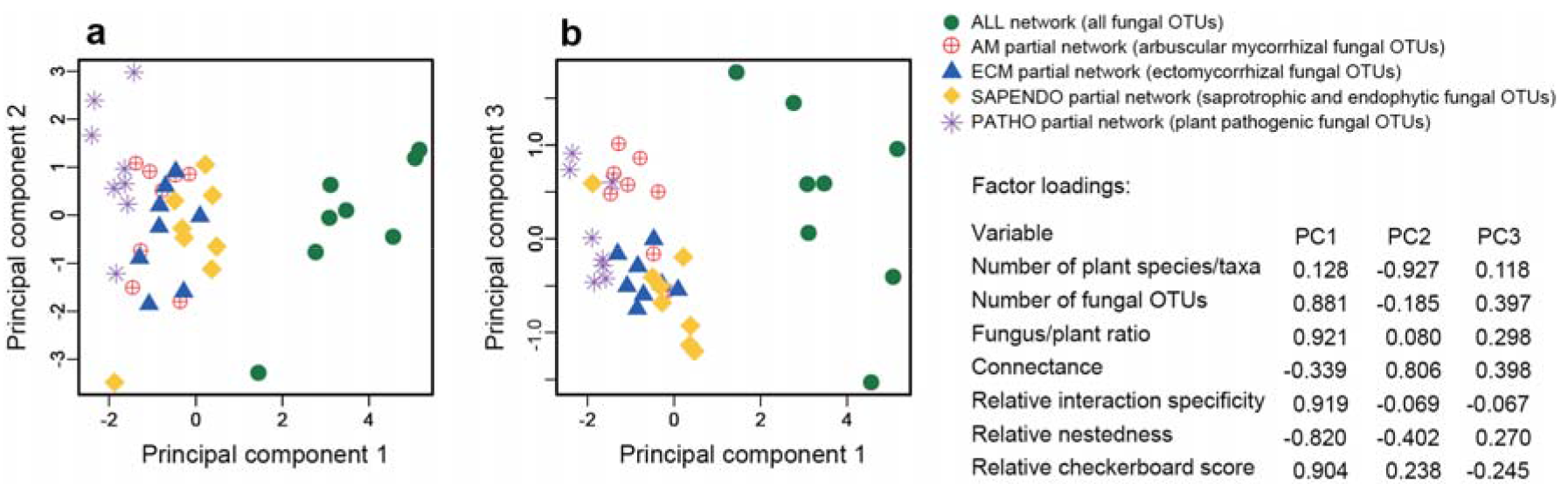
Principal component analysis of network properties. **a** Principal component 1 vs. principal component 2. Factor loadings of the examined variables are shown on the right. **b** Principal component 1 vs. principal component 3.

## Discussion

Our data, which included 17–55 plant species/taxa and more than 1000 fungal OTUs in each of the eight forests, provided a novel opportunity to evaluate how different types of below-ground plant–fungus associations varied in their community-scale characteristics along a latitudinal gradient. We then found that network structural properties differed among different types of plant–fungus associations (Fig. 3), while geographic factors contributed to the variation found in network structure (Table 1). Specifically, arbuscular mycorrhizal networks differed in their architecture from ectomycorrhizal and saprotrophic/endophytic networks, characterized by high connectance (Fig. 4). We also found that networks consisting of all functional groups of fungi and their host plants had higher network-level interaction specificity, more differentiated host ranges between fungi, and lower network nestedness than the partial networks of arbuscular mycorrhizal, ectomycorrhizal, and saprotrophic/endophytic associations (Figs. 3 and 4). As in previous studies, our data included many fungal OTUs unassigned to functional groups due to the paucity of the information of fungal functions and guilds in databases [56]. However, by extending findings in previous plant–fungus network studies [30, 31, 65], in which sampling strategies, interaction type, or geographic factors were not controlled systematically, this study offers a basis for discussing how different types of below-ground plant–fungus associations collectively build plant–soil feedback in terrestrial ecosystems.

Among the network indices examined in this study, nestedness showed an idiosyncratic tendency in light of other types of interaction networks examined in community ecology [34–36]. We found that below-ground plant–fungus networks often displayed “anti-nested” architecture, in which scores representing nested network structure were lower than those expected by chance (i.e., negative values of relative nestedness; Fig. 3e), as suggested also in previous studies [17, 31, 65]. Although factors organizing anti-nested network architecture remain to be investigated, competition for host plants among fungal species has been inferred to decrease nestedness of plant–fungus associations [65]. In addition, a previous comparative study suggested that plant–fungus network nestedness decreased with increasing annual mean temperature on a global scale [27].

The prevalence of anti-nested or non-nested network structures is in sharp contrast to observations on other types of plant–partner networks, which commonly show statistically significant nested architecture [34]. Specifically, plant–pollinator and plant–seed disperser interactions are generally characterized by nested network architecture in which overlap of partner ranges within the same guild are expected to mitigate competition between plant species [34–36]. Therefore, anti-nested structure of plant–fungus networks provides important insights in terms of community ecological theory linking network structure and species coexistence [36, 37, 70]. Given that below-ground fungi constitute one of the most species-rich components of the terrestrial biosphere [3], understanding community-scale properties of below-ground plant–fungus associations is a major step for disentangling relationship among network structure, species coexistence, and community stability.

To overcome the inconsistency between theory and observations, we may need to take into account basic biology of below-ground plant–fungus associations. We here highlight two backgrounds that need more attention for deepening discussion on ecological networks and species coexistence. First, in contrast to plant–pollinator or plant–seed disperser networks, which are often assumed to consist only of mutualistic interactions, below-ground plant–fungus networks can involve not only mutualistic but also antagonistic and commensalistic interactions. This diversity of interaction type can lead to high stability of below-ground fungal and their host plant communities. Specifically, while communities consisting exclusively of mutualistic interactions are inherently unstable [71], involvement of a small fraction of antagonistic interactions in those communities can dramatically enhance species coexistence [72]. Second, because fungi can disperse long distances as spores [73, 74] (but see [75]), their local species richness (alpha diversity) may be greatly impacted by metacommunity processes [76]. Interestingly, a recent theoretical study on food webs predicted that strong coupling of local communities within a metacommunity could result in positive relationship between species richness and community stability [77]. Such theoretical evaluation of metacommunity dynamics has been extended to systems involving mutualistic interactions [78], providing platforms for considering how dispersal abilities of constituent species determine local species richness/coexistence of different types of plant–partner networks.

To apply a standardized criterion for plant–fungus associations, we did not perform any data screening based on the “mycorrhizal types” of plant species. As a result, our data included plant–fungus combinations that could not be classified into well-recognized categories of mycorrhizal symbioses [8]. For example, ectomycorrhizal fungi were detected not only from plant species in “ectomycorrhizal” families (e.g., Fagaceae, Pinaceae, and Betulaceae) but also from other plant species (Fig. 2; Additional file 6: Data S6). In addition, the data included network links between arbuscular mycorrhizal fungi and ectomycorrhizal plant species (Additional file 6: Data S6) as reported previously [79]. Although such plant–fungus associations that do not fall into classic categories of mycorrhizal symbioses seldom attract attention and they are often removed from high-throughput sequencing datasets, some of them may represent important ecological interactions. An ectomycorrhizal fungus in the truffle genus (*Tuber melanosporum*), for instance, is known to cause severe necrosis in root cortices of non-ectomycorrhizal herbaceous plants [80]. Thus, for the standardization of plant–fungus network analyses inferred with high-throughput sequencing, we need to take into account the possibility that network links can represent not only mutualistic but also neutral and antagonistic interactions [17]. Given also that even well-known combinations of plant–fungus mycorrhizal interactions can result in antagonistic interaction depending on soil environmental conditions and host plant nutrition [81, 82], potential diversity of ecological interactions within a network and its community-scale consequences [72] deserve intensive research.

Our community-scale comparative analysis targeting a broad latitudinal range from cool-temperate to subtropical regions has some implications for geographic diversity patterns of plant–associated fungi, although careful interpretation is required given the small number of study sites. The number of detected ectomycorrhizal fungal OTUs was lower in subtropical than in temperate forests (Fig. 3a), presumably reflecting geographic variation in the relative abundance of Fagaceae, Pinaceae, and Betulaceae plants in plant communities as discussed in previous studies [83–86] (see also [87]). In contrast, the number of arbuscular mycorrhizal fungal OTUs increased towards south in our data, while a previous meta-analysis detected no latitudinal diversity gradient regarding the fungal functional group [88] (see also [89]). The total number of fungal OTUs was also higher in subtropical forests, peaked in the southernmost site. Interestingly, unlike other study sites, the southernmost sampling site was characterized by low levels of network-scale interaction specificity and host plant differentiation as well as by the absence of anti-nested network architecture. Although some pioneering studies have investigated host preferences of tropical fungi [90–92], it remains a major challenge to examine whether the observed latitudinal gradient in plant–fungus network structure is extended to tropical regions.

## Conclusions

Based on the large datasets of root-associated fungi, we herein showed how plant–fungus network architecture vary across a latitudinal gradient across the Japanese Archipelago. For further understanding the diversity of below-ground pant-fungus associations, more comparative studies of community-scale characteristics are required especially in the tropics. Moreover, further data of networks consisting of pathogenic fungi and their host plants are awaited to discuss community-scale properties of negative plant–soil feedbacks [93]. Given that the number of pathogenic fungi included in our present analysis was too few to evaluate statistical features of their networks, selective sampling of pathogen infected plant individuals may be necessary. Improving reference databases of fungal functions is also an important challenge towards better understanding of the roles of fungal communities. More macroecological studies of plant–fungus interactions [75, 94, 95], along with experimental studies testing functions of poorly characterized fungi [11, 13, 14], will reorganize our knowledge of terrestrial ecosystem processes.

## Abbreviations

ANOVA: analysis of variance
DDBJ: DNA Data Bank of Japan
ITS: internal transcribed spacer
LCA: lowest common ancestor
OTU: Operational taxonomic unit
QCauto method: query-centric auto-*k*-nearest neighbor method

## Acknowledgements

We thank Teshio Experimental Forest (Hokkaido University), Tomakomai Experimental Forest (Hokkaido University), Sugadaira Research Station (Tsukuba University), Yona Field (Ryukyu University), Tropical Biosphere Research Center (Ryukyu University), and Forestry Agency of Japan for the permission of fieldwork and Takayuki Ohgue, Takahiko Koizumi, Miyako Natsume, Ayu Narita, Yoriko Sugiyama, and Yamato Unnno, Yuko Sawanobori, and Minato Kodama for their support in molecular experiments.

## Availability of data and materials

The Illumina sequencing data were deposited to DNA Data Bank of Japan (DDBJ Sequence Read Archive: DRA006339). The raw data of fungal community structure and the fungal community matrices analyzed are available as Additional files 1-6.

## Authors’ contributions

HT designed the work. HT, HS, SY, and AST conducted fieldwork. HT, HS, and SY performed the molecular experiments. HT wrote the manuscript with HS, SY, and AST.

## Consent for publication

Not applicable

## Ethics approval and consent to participate

Not applicable

## Additional files

**Additional file 1: Data S1.** Information of study sites, taxonomic and functional-group information of the fungal OTUs detected.

**Additional file 2: Data S2.** Sample-level matrices of plant–fungus associations.

**Additional file 3: Data S3.** Sequences of the non-glomeromycete fungal OTUs detected.

**Additional file 4: Data S4.** Sequences of the glomeromycete fungal OTUs detected.

**Additional file 5: Data S5.** Species-level matrices of plant–fungus associations.

**Additional file 6: Data S6.** Network data matrices.

**Additional file 7: Data S7.** Results of the randomization analysis.

